# Advancing Reef Monitoring Techniques through Exometabolomics: Quantification of Labile Dissolved Organic Metabolites on Coral Reefs

**DOI:** 10.1101/2023.12.20.572630

**Authors:** Brianna M. Garcia, Cynthia C. Becker, Laura Weber, Gretchen J. Swarr, Melissa C. Kido Soule, Amy Apprill, Elizabeth B. Kujawinski

**Author notes:** **Corresponding Authors** Amy Apprill - *Department of Marine Chemistry and Geochemistry, Woods Hole Oceanographic Institution, Woods Hole, MA, United States*, Elizabeth B. Kujawinski - *Department of Marine Chemistry and Geochemistry, Woods Hole Oceanographic Institution, Woods Hole, MA, United States. Department of Soil, Water, and Ecosystem Sciences, University of Florida, Gainesville, FL, United States.

## Abstract

Extracellular chemical cues constitute much of the language of life among marine organisms from microbes to mammals. These chemical cues shape foraging and feeding strategies, symbiotic interactions, selection of mates and habitats, and the transfer of energy and nutrients within and among ecosystems. Changes in the chemical pool flux within these ecosystems serve as invisible signals of overall ecosystem health and, conversely, disruption to this finely tuned equilibrium. In coral reef systems, the full scope and magnitude of the chemicals involved in maintaining reef equilibria are largely unknown. In particular, processes involving small, polar molecules, which form the majority components of labile dissolved organic carbon (DOC), are often poorly captured using traditional techniques and therefore remain unconstrained. In this study, we employed a recently developed chemical derivatization method and mass spectrometry-based targeted exometabolomics to capture and quantify polar dissolved phase metabolites on five coral reefs in the U.S. Virgin Islands. We detected and quantified 45 polar exometabolites, and further demonstrate their variability across the investigated geographic landscape and contextualize these findings in terms of geographic and benthic cover differences. By comparing our results to previously published coral reef exometabolomes that did not employ chemical derivatization, we show the novel quantification of 23 metabolites which were previously undetected in the coral reef exometabolome, including compounds involved in central carbon metabolism (*e.g.,* glutamate) and novel metabolites such as homoserine betaine. We highlight the immense potential of chemical derivatization-based exometabolomics for quantifying labile chemical cues on coral reefs and measuring molecular level responses to environmental stressors. Overall, improving our understanding of the composition and dynamics of reef exometabolites is vital for effective ecosystem monitoring and management strategies.

**Synopsis:** Minimal research exists on the polar exometabolites on coral reefs, an essential component of reef health and the majority component of labile DOC. This study reports 23 newly quantified polar metabolites on coral reefs and highlights significant differences in the exometabolome composition across geographically distinct reefs and bays.

## Introduction

The immense diversity of small organic molecules released and transformed by organisms in aquatic environments has long been a barrier to understanding organismal interactions and ecosystem processes.^1^ Recent methodological and technological advances in chemistry (*e.g.,* better separation methodologies, sensitivity, accuracy, and resolving power) have made the detection and quantification of low-concentration, high-flux compounds more tractable.^2,3^ Because of its unmatched sensitivity, mass spectrometry (MS) has become the dominant and most widely used technology for the chemical characterization of marine dissolved organic matter (DOM). However, directly analyzing low-concentration organic compounds in seawater containing high concentrations of salt is extremely challenging. For example, sulfate and chloride ions comprise tens to hundreds of millimolar (mM) concentrations in seawater, respectively. Comparatively, steady-state concentrations of bulk dissolved organic carbon (DOC) occur in the micromolar (□M) range, with individual compounds present at nanomolar (nM) to picomolar (pM) concentrations.^4^

Given the limitations of direct analysis, various methods have been developed to concentrate and isolate marine DOM including tangential flow filtration (TFF),^5^ reverse osmosis/dialysis (RO/ED),^6^ and solid-phase extraction (SPE).^7^ However, these approaches result in significant biases in the chemical species retained for analysis. In particular, small and polar metabolites (characteristics of many labile biomolecules and majority drivers of carbon flux) are lost during TFF or RO/ED and are not well retained (extraction efficiencies < 50%) on SPE resins such as the commonly used styrene-divinylbenzene polymer (PPL) resin.^8^ While a subset of these polar metabolites have been detected using SPE,^8^ metabolites with low standing stock concentrations, quick turnover times, or highly polar moieties are largely overlooked. To overcome some of these limitations, alternative sample preparation approaches have been developed allowing for specific chemical species to be retained, concentrated, and analyzed. Specifically, alternative SPE resins (*i.e.,* cation exchange^9^) and chemical derivatization ^10–12^ have recently been shown to improve the analysis of otherwise elusive polar compounds. While the development of these methodologies has been described, their widespread application to various marine ecosystems remains limited. Further employment of these methodological advancements will allow for the identification and quantification of a wider range of the labile dissolved molecules that are rapidly consumed by heterotrophic bacteria and other organisms, thus enhancing our baseline understanding of these systems.^4^ By expanding our view of the chemical “players’’, we can begin to decipher the complex relationships between sources and sinks of metabolites and how they change over time (flux), under stress (*e.g.,* incidents of disease), and with global climate changes.

One such environment that would benefit from these innovative methods are coral reefs. Coral reefs are one of the most productive, biologically diverse, and economically valuable ecosystems worldwide.^13^ Yet, our understanding of the chemistry occurring within reef systems remains limited, with large knowledge gaps in chemical composition, biogeochemical cycles, and chemical diversity (regionally and in the face of global changes/stressors). Previous studies have shown that extracellular DOM released in a species-specific matter by benthic primary producers plays a key role in fueling the reef food web and regulating ecosystem function.^4,14,15^ Unfortunately, coral ecosystems have undergone significant decline due to climate change, natural disturbances, disease, and anthropogenic factors, resulting in changes in benthic and microbial composition.^16^ DOM released by altered communities of benthic primary producers may thereby negatively impact coral health by changing microbial community structure, perhaps favoring coral pathogens, and/or increasing microbial respiration causing localized hypoxia.^14,17^ Such findings underscore the need to transition from bulk measures of DOM to analyses of individual molecules in order to better understand changes in the composition of reef DOM.

Metabolomics studies using untargeted approaches have been more widely, yet still modestly, used to generate a molecularly resolved view of the total DOM pool in reef seawater and tissue homogenate. However, targeted approaches employing matched internal standards are required to determine absolute quantitation and confident metabolite identifications. Metabolomics measurements of dissolved exudates from coral reefs using the conventional PPL-SPE approach (both targeted and untargeted) have investigated a range of coral reef phenomena including metabolites associated with pristine reef systems,^18,19^ non-self interactions,^19^ bleaching,^20^ disease,^21^ and benthic-pelagic coupling.^22^ However, our knowledge on these topics is still limited because these datasets lack detection of some of the most polar compounds, which are likely central components of nutrient and energy cycling on oligotrophic coral reefs.

Herein, we identify and quantify the exometabolome of five coral reefs in St. John, U.S. Virgin Islands located within the Virgin Islands National Park using the pre-extraction benzoyl-chloride derivatization method developed by Widner *et al.*^10^ and targeted liquid chromatography-tandem mass spectrometry. We compare our findings to previously quantified reef exometabolomes and describe 23 newly quantified compounds on coral reefs. Additionally, we compare the polar exometabolomes over five geographically separated reefs and describe trends related to benthic composition and potential anthropogenic influences. This dataset significantly expands our lexicon of polar, labile metabolites present on coral reefs.

## Material and methods

### Chemicals and standards

Hydrochloric acid (HCl), acetone, methanol (MeOH), acetonitrile (MeCN), benzoyl chloride (BC; 99%, ACROS Organics), and sodium hydroxide (NaOH) were purchased from Fisher Scientific (Optima). Phosphoric acid (85%, ACS reagent grade), ^13^C_6_-ring-BC (99% atom ^13^C), and internal standards were purchased from Sigma-Aldrich. Stable isotopically labeled - internal standards matched to each targeted compound (SIL-IS) were prepared with ^13^C_6_-ring-BC. Deionized water was obtained from a Milli-Q system (Millipore; resistivity 18.2 MΩ at 24 °C, TOC < 1 μM). Samples and reagents were stored in acid-washed, combusted (at least 4 h at 450°C) glassware. BC and all solvents were transferred using combusted glass Pasteur pipettes. Primary stocks and mixes were stored at −20°C. The working reagent (5% BC in acetone) was prepared fresh daily.

### Sample collection

Samples were collected in January 2021 in St. John, U.S. Virgin Islands within the Virgin Islands National Park (Figure 1; Table S1). Seawater was collected at five coral reefs, two located in Lameshur Bay: Tektite (TK, n=3 18.309°N, 64.723°W, 8.3 m reef depth) and Yawzi (YZ, n=3, 18.314°N, 64.726°W, 8.4 m), and three in Fish Bay: Ditliff (DL, n=4, 18.313°N, 64.764°W, 6.10 m), Cocoloba (CO, n=4, 18.315°N, 64.761°W, 7.01 m), and Joel’s Shoal (JS, n=3, 18.313°N, 64.757°W, 8.8 m). All reefs fell within the Virgin Islands National Park except for DL, which is on the eastern edge.

**FIGURE 1.**
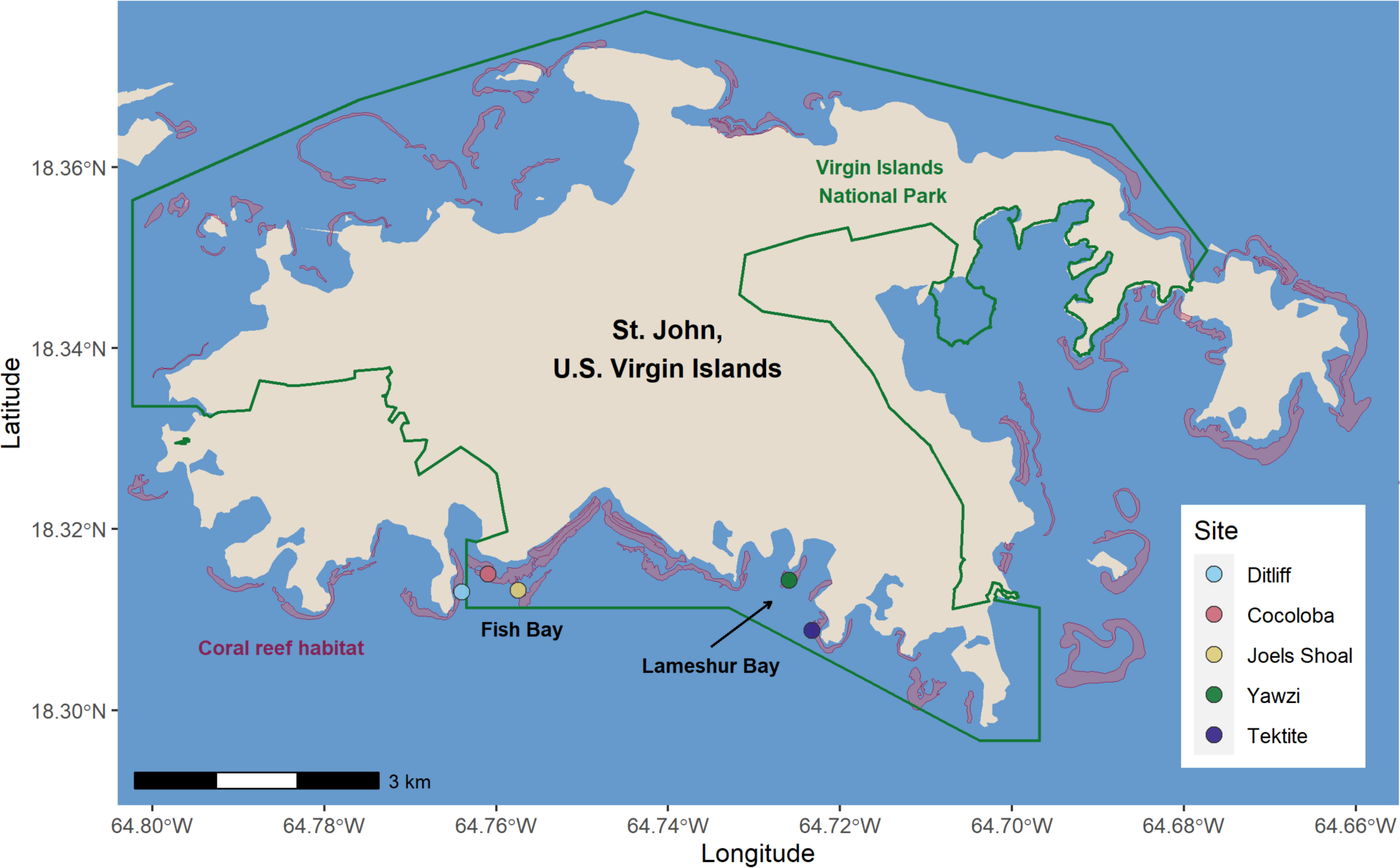
Map of St. John, USVI (tan) and the Virgin Islands National Park (green boundary) Coral reefs are depicted around the island (pink areas). Coral reefs where reef water was collected are shown using circles and labeled by reef name in the legend. The two bays where reefs were located are shown labeled on the map.

Scuba divers collected reef depth seawater (within 0.25 m of benthos) in acid-washed Niskin Bottles (General Oceanics, Miami, Florida, United States). Specifically, divers descended with three Niskin bottles in the open position, thoroughly rinsed the Niskin with reef depth seawater, located three areas of the reef that were topographically complex, and triggered each Niskin to close, capturing reef depth seawater within the Niskin chamber at three distinct locations on each reef. After ascent, the Niskins were drained into bottles, placed on ice and processed within 2 hours of collection.

### Reef and Microbial Descriptions

#### Benthic Survey

Point intercept benthic surveys using 10 m transects were conducted at four random locations at each reef during January 2021. Every 10 cm, the underlying biological reef organisms or substrate was recorded with the following categories: hard coral, macroalgae, cyanobacterial mats (CYAN), crustose coralline algae (CCA), diseased coral, the invasive crustose coralline algae *Ramicrusta*, sponge, soft coral, turf algae, substrate (non-biological), and other. The “other” category included other invertebrates, *Millepora*, dead coral, and eelgrass. Counts of individual categories were converted to relative abundance. The benthic cover across reefs was compared with a principal component analysis using the PCA function from FactoMineR (v2.6) R package^23^ with each value scaled to unit variance, followed by visualization with the *fviz_pca* function from the factoextra (v1.0.7) R package.

#### Dissolved Organic Carbon and Microbial Abundances

From each Niskin water collection described above, 40 mL of reef depth seawater was processed and analyzed for non-purgeable organic carbon (NPOC) and total nitrogen (TN) and 1.4 mL was collected, processed, and analyzed for the enumeration of *Prochlorococcus*, *Synechococcus*, picoeukaryotic cells, and unpigmented cells (heterotrophic bacteria and archaea) following methods described in the Supplemental Methods and *Weber et al. 2022*^22^, respectively.

### Metabolite Processing

#### Metabolomics

For metabolomics analysis (Figure 2), 10 L of reef depth seawater collected within the individual Niskin bottles were transferred into multiple acid-washed 1 or 2 L polycarbonate bottles using acid-washed thermoplastic elastomer (TPE) tubing (PharMedBPT MasterflexTM, Cole – Parmer, Vernon Hills, IL, United States). Seawater was filtered through polytetrafluoroethylene (PTFE) 0.2 μm pore size, 47 mm filters (Omnipore, EMD Millipore Corporation, Billerica, MA, United States) using peristalsis (MasterFlex L/S pump and pump heads, Cole-Parmer, Vernon Hills, IL, United States). TPE tubing and acid-washed fluorinated ethylene propylene (FEP) tubing (890 Tubing, Nalgene^TM^, Thermo Scientific^TM^, Waltham, MA, United States) were used to transfer pumped seawater through the filter membrane and into acid-washed polycarbonate collection bottles. Subsamples of filtrate for each sample were added into combusted 40 mL amber vials for benzoyl-chloride derivatization. Samples were stored and transported at −20°C until sample processing and derivatization was conducted.

**FIGURE 2.**
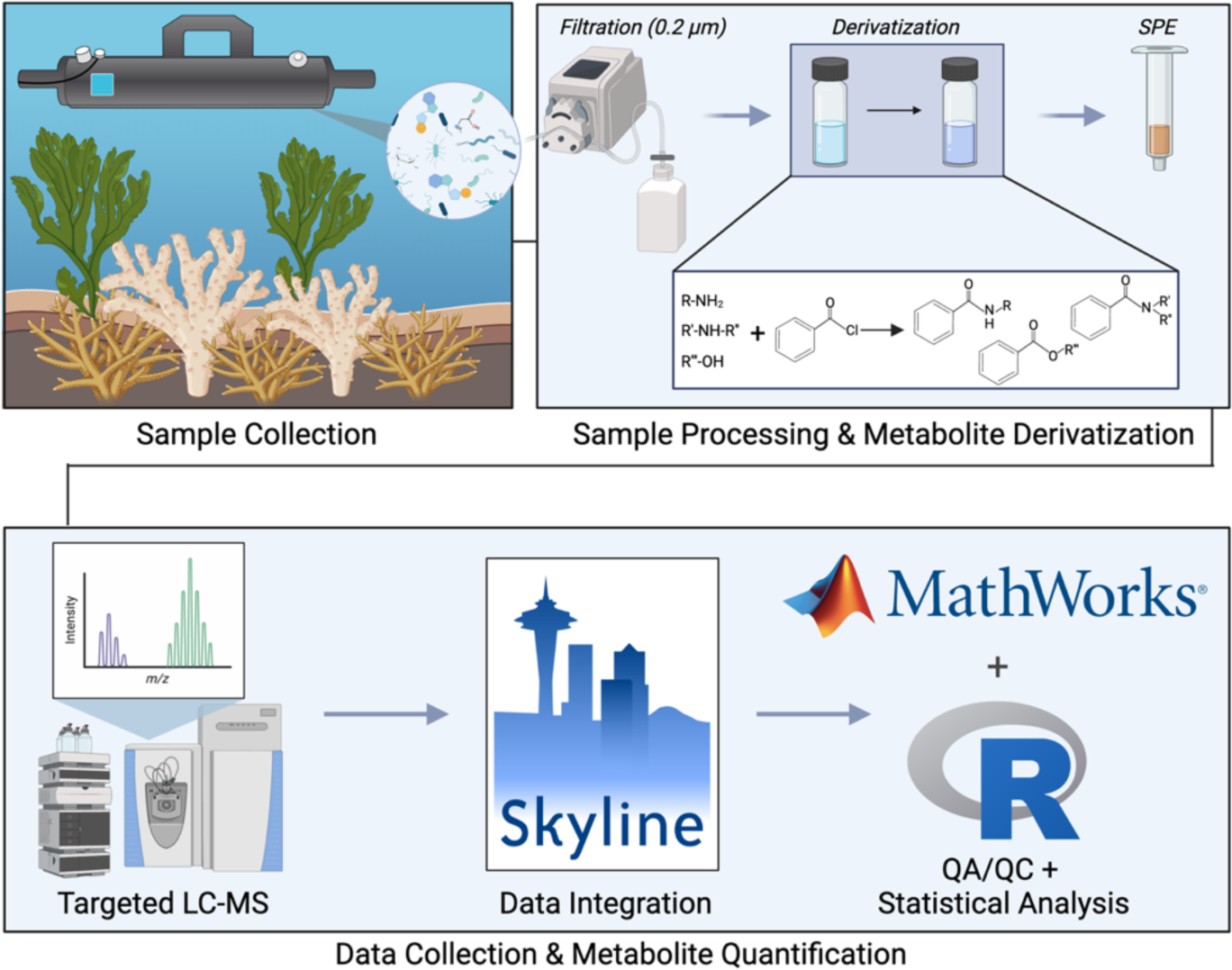
Experimental design for metabolomics analysis of coral reef exometabolites and benzoyl-chloride derivatization. Sample Processing & Metabolite Derivatization panel modified from Widner *et al.* 2021. R logo © 2016 The R Foundation, CC-BY-SA 4.0.

#### Benzoyl-chloride metabolite derivatization and extraction

Filtered, frozen, seawater samples were thawed and 25 mL from each sample was transferred into a fresh amber glass vial for ^12^C-benzoyl chloride derivatization conducted according to Widner *et al.*^10,24^ A standard curve was derivatized (^12^C-BC) in parallel with samples, using off-reef seawater as the matrix. The standards spanned a concentration range of 0-25 ng added of each standard. ^13^C-labeled stable isotopically labeled – internal standards (SIL-IS) were similarly derivatized in parallel, using ^13^C-benzoyl chloride, and spiked into all experimental and standard curve samples. Derivatization methods are described in detail in the Supplemental Methods. Derivatized samples were then extracted by SPE using 6 mL, 1 g Bond Elut PPL cartridges (Agilent, Santa Clara, CA, USA), and the final eluent dried via vacufuge to near-dryness at 30°C. Dried samples were stored at −20°C until LC-MS analysis, at which time each sample was reconstituted in 5% MeCN, transferred to a 2 mL LC vial (with small volume insert), and topped with 5 μL of 100% MeCN. Samples were stored at 4°C until instrumental analysis.

### Targeted metabolomics UHPLC-MS/MS data collection

Reverse phase (RP) chromatography was performed according Widner *et al*.^10^ Using an ultra-high performance liquid chromatography system (UHPLC; Vanquish, Thermo Scientific, Waltham, MA, USA), separation was achieved on a Waters Acquity HSS T3 column (100Å, 1.8 µm particle size, 2.1 mm X 100 mm) equipped with an Acquity HSS T3 VanGuard pre-column (100Å, 1.8 µm particle size, 2.1 mm X 5 mm) coupled to a heated electrospray ionization source (H-ESI) and an Orbitrap mass spectrometer (Orbitrap Fusion Lumos, Thermo Scientific). Chromatography gradient, mass spectrometer settings, and quality assurance and control (QA/QC) are described in detail in the Supplemental Methods.

### Metabolomics data processing

Thermo .raw files were processed using Skyline (v. 21.2.0.568), an open-source and freely available software package for targeted small molecule quantification.^25,26^ Metabolites were detected and peaks integrated based on accurate mass (within a 10 ppm mass tolerance), retention time (RT), and MS/MS fragment ion confirmation. Calibration curves for each compound (9 points each) were constructed based on the amount of metabolite standard added (ng added) versus the integrated light-to-heavy peak area ratios calculated using Skyline. Exported quantification tables from Skyline containing peak areas for each metabolite were imported into MATLAB^®^ (v. R2022a). An in-house MATLAB^®^ script (*considerSkyline.m*) was used to generate linear regression calibration curves, quantify each metabolite, calculate limits of detection (LOD) and quantification (LOQ), and merge positive and negative polarity mode data (further described in Supplemental Methods). All MATLAB^®^ scripts used for processing the Skyline outputs are available online at https://github.com/KujawinskiLaboratory/SkyMat. The resulting merged quantification table (SI Excel Table S1) was further filtered in MATLAB^®^ based on the QA/QC metrics described in detail in the Supplemental Methods.

### Metabolite Comparison Across Studies

Comparisons between the metabolites observed in this study using the BC derivatization method and four independent, targeted benthic exometabolomics datasets that did not employ BC derivatization were investigated. Studies were selected due to their quantitative measurements of exometabolites on coral reefs allowing for valid comparisons, and included reef seawater samples collected in Cuba^18^ and along the coast of Florida,^27^ a U.S. Virgin Islands coral reef seawater incubation experiment,^22^ and Florida field-collected sponge inhalant and exhalant seawater samples.^28^ All four studies utilized PPL SPE extraction (without derivatization), the most common methodology employed for characterizing DOM by LC-MS described by Dittmar *et al.*^7^ Metabolite names from each study were downloaded from each manuscript’s publicly available metadata. Metabolites described as unresolved (*i.e.,* “homoserine and threonine” in Weber *et al.* 2020) were excluded from the analysis. Metabolite names were converted to InChIKeys using the Chemical Translation Service (SI Excel Table S2).^29^ In instances where no conversion was possible, InChIKey were obtained manually from the Public Chemical Database (PubChem).^30^ The International Chemical Identifier code (InChI) or InChIKey annotates metabolites by their chemical structures allowing for more standardized comparisons.^31^ The collective metabolite names and their corresponding InChIKeys from all five studies were imported into R. Unique InChIKeys were queried across all studies producing a binary matrix where a value of one indicated the presence of that metabolite in a study, and zero indicated the metabolite’s absence (SI Excel Table S2). The UpSetR package *upset* function was used to generate an upset plot depicting the overlap of metabolites across studies.^32^

### Statistical Analysis

Statistical analyses were carried out in R Studio (v. 2022.12.0 Build 353) running R (v4.2.2, 2022-10-31).^33,34^ All code and data used for recreating figures are publicly available on GitHub (https://github.com/bmgarcia/CINAR_Habitat_Metabolomics). Data was tested for normality visually via histogram plots and mathematically using the Shapiro–Wilk test (*shapiro.test*). The majority of metabolites were not normally distributed (P < 0.05) and thus non-parametric tests were used for all downstream analyses. Non-metric multidimensional scaling (NMDS) using the ‘vegan’^35^ and ‘ggplot2’^36^ packages was used to visualize the Bray Curtis dissimilarity between metabolomics samples. A Permutational Multivariate Analysis of Variance (PERMANOVA) using Bray Curtis distance matrices was carried out on metabolite and benthic data using the *adonis2* function in the ‘vegan’ package. A Kruskal-Wallis test (*kruskal.test*) was used to test for significant metabolites between reefs and bays (significance level P ≤ 0.05). Pairwise Wilcoxon Rank Sum tests (*pairwise.wilcox.test*) were conducted on significant metabolites from the Kruskal-Wallis test to determine pairwise significance. P-values were adjusted to account for multiple comparisons using the Benjamini-Hochberg correction^37^ and metabolites were considered significant with a p_adj_ value ≤ 0.1. Pairwise Spearman correlation coefficients were determined between metabolite concentrations and benthic survey proportions using the *cor* function in the base ‘stats’ R package (v4.2.2.). Statistical results can be found in the Supplemental Information (SI Excel Table S3-4).

## Results and Discussion

### Metabolites released from coral reefs

Using pre-extraction benzoyl-chloride derivatization coupled with UHPLC-MS/MS targeted metabolomics on reef water collected above five coral reefs in the U.S. Virgin Islands, we detected 45 dissolved phase metabolites out of the 79 metabolites in this method. The number of detected metabolites varied by reef; 31 metabolites were present at all five reefs and 14 fell below the limit of detection at one or more locations (Figure 3; Table S1). Exudates from JS had the highest number of detected metabolites (44), followed by TK and YZ (39), and DL and CO (35). Of the 45 metabolites, two were detected and 43 were quantified (SI Excel Table S5-quantified is defined herein as a metabolite having a measured value greater than the limit of quantification (LOQ) in at least one sample). Metabolite concentrations ranged from 5 pM to 52 nM with a median concentration of 390 pM (Figure 3). Of the 45 metabolites that were captured using the BC method, 30 had reported extraction efficiencies (EE) of less than 1% using PPL SPE (without derivatization), highlighting the longstanding challenges of detecting these polar metabolites in seawater.^8^ For example, DHPS (2,3-dihydroxypropane-1-sulfonate) which is poorly retained using PPL SPE (EE < 1%), was common across all reefs at relatively high concentrations ranging from 600 pM to 52 nM, with a median concentration of 2 nM. DHPS is an organosulfur molecule hypothesized to play a major role in the flux of sulfur and carbon through marine food webs.^38,39^ Reef-building corals are holobionts comprised of the cnidarian host and their associated microbes including dinoflagellates and other protists, bacteria, archaea, fungi, and viruses. Significant concentrations of other sulfur-containing metabolites (*i.e.,* dimethylsulfoniopropionate [DMSP] and dimethyl sulfide [DMS]) have been recorded in coral reef hosts and their symbiotic dinoflagellates, suggesting that coral reefs might play a substantial role in sulfur cycling in largely oligotrophic regions.^40–43^ Unlike DMSP, our understanding of the breadth of DHPS producers and their role in sulfur biogeochemistry remains limited. Sulfonates such as DHPS are emerging as important chemical links between marine phytoplankton and bacteria, which together fuel the microbial sulfur cycle in the surface ocean.^44,45^ The identification of DHPS on coral reefs in this study requires further investigation into its source(s) (*i.e.,* coral host, *Symbiodinium spp.*, or associated microbes) and its role(s) as a potential key player in sulfur cycling in coral reefs. The use of alternative chemical approaches, such as chemical derivatization, are key to increasing our understanding of these important labile biomolecules in the context of marine microbial systems.

**FIGURE 3.**
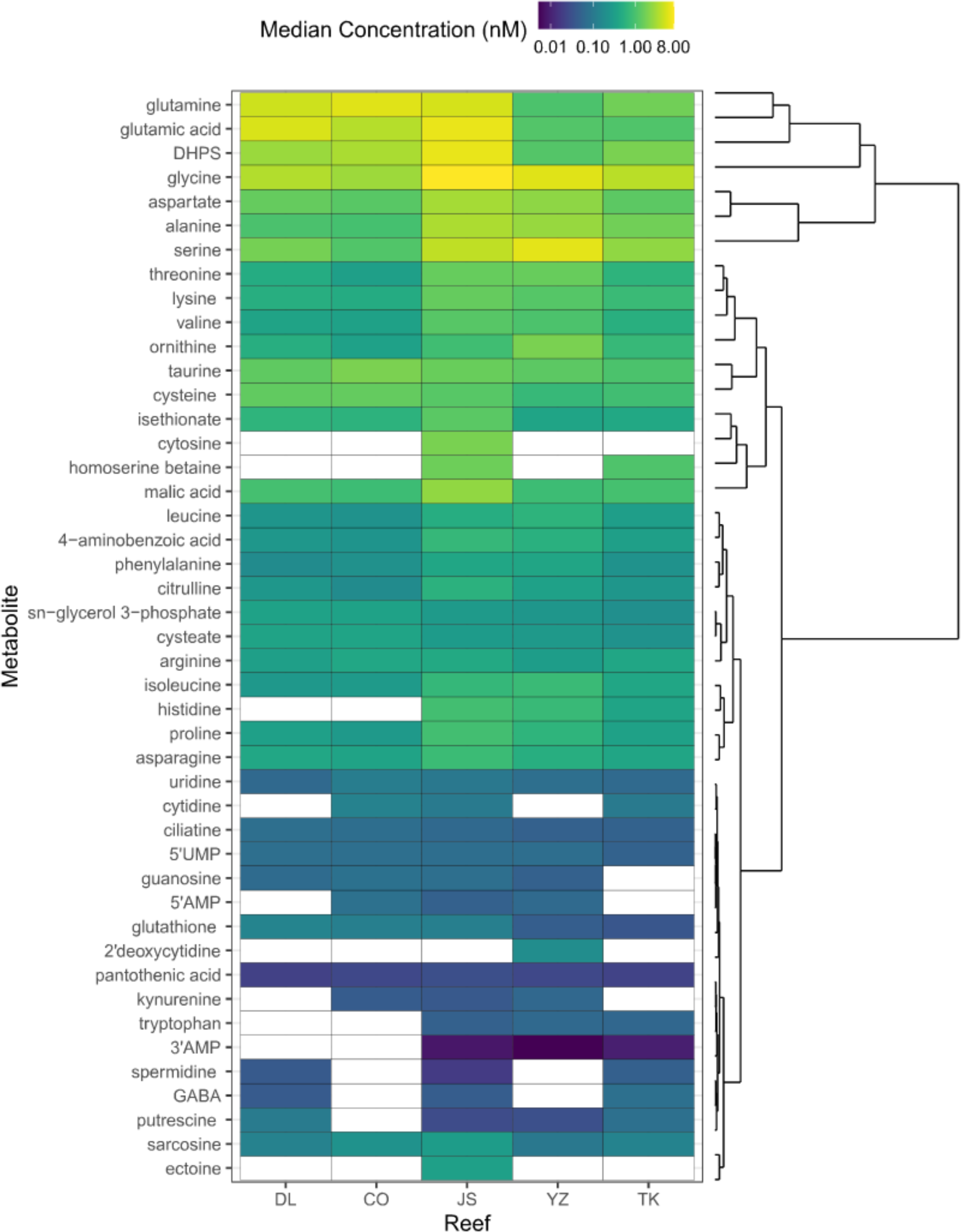
Metabolites were common across five coral reefs. Heatmap of log-transformed dissolved metabolite concentrations (nM). Values shown are the median concentration across replicates (n=3-4). Warm colors reflect higher concentrations whereas cooler colors represent lower concentrations. Concentrations that fell below the LOD in all replicates are depicted using a white fill. X-axis labels reflect the five coral reefs (DL - Ditliff, CO - Cocoloba, JS - Joel’s Shoal, YZ - Yawzi, and TK - Tektite), and y-axis labels indicate the metabolite name. Metabolites were clustered based on metabolite concentrations and ordered accordingly. Hierarchical clustering is shown as a dendrogram adjacent to the y-axis.

### Benzoyl-chloride derivatization retains polar metabolites previously undetected in the coral reef exometabolome

To determine the advantages and complementarity of incorporating pre-extraction benzoyl-chloride derivatization into the marine metabolomics workflow, we assessed the overlap of identified metabolites across coral reef benthic communities from four previously published PPL-SPE extracted reef water targeted metabolomics datasets (Fiore *et al.* 2017,^28^ Weber *et al.* 2020,^18^ Weber *et al.* 2022,^22^ and Becker *et al.* 2023^27^) and the current study (Figure 4). A total of 85 distinct metabolites were identified across the five studies. Of these 85 metabolites, 38 were found to be study-dependent (only identified in a single study), 41 were identified in a minimum of two studies, and six were ubiquitously identified (Figure S1). The largest number of identified metabolites (53) was by Weber *et al.* 2020 on highly protected and biodiverse Cuban coral reefs followed by the current study’s benzoyl-chloride derivatized dataset (45). Variations observed in metabolite overlap are unlikely to be platform dependent since all datasets were collected on the same mass spectrometry platform.

**FIGURE 4.**
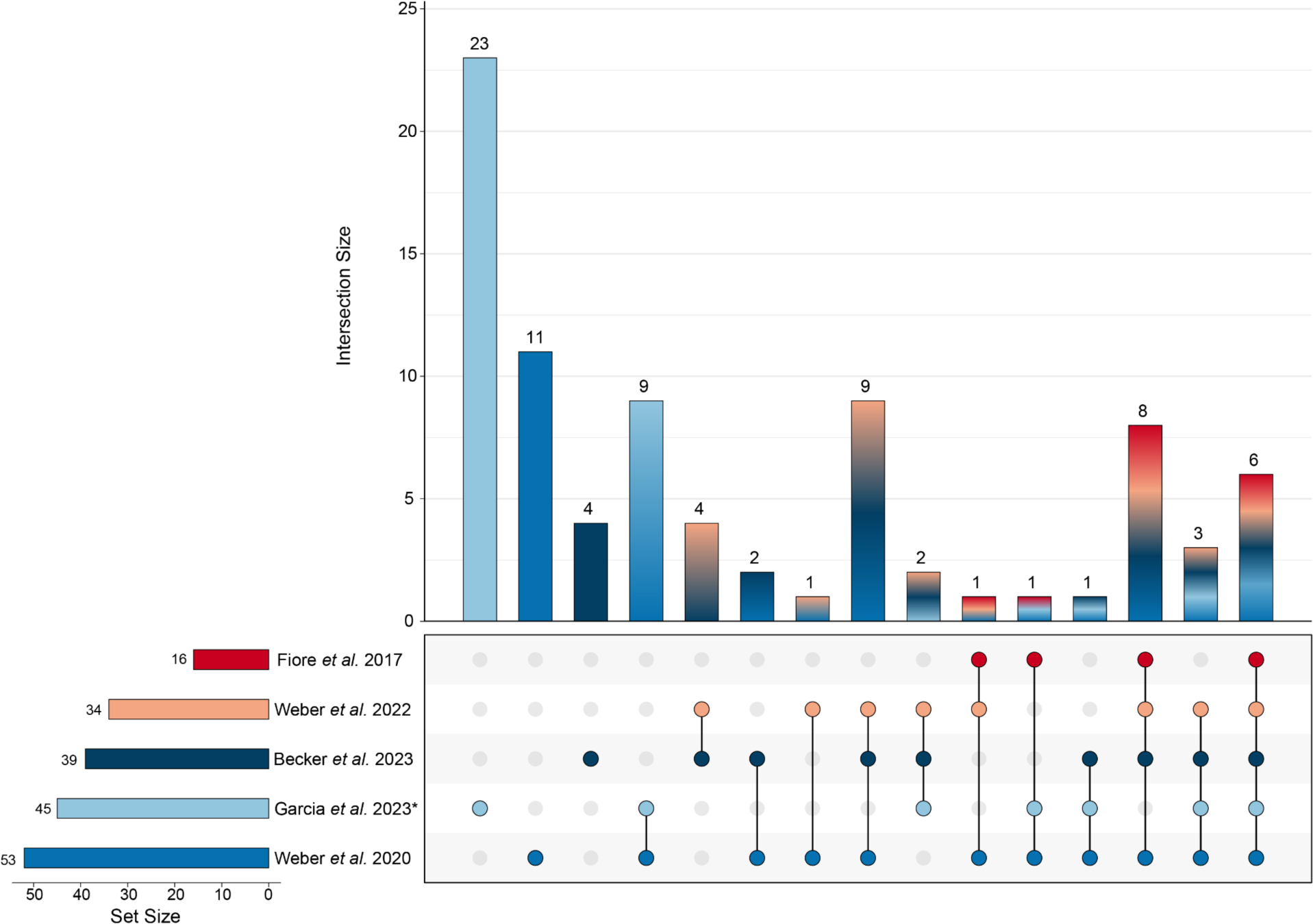
Comparison of coral reef studies shows that 23 metabolites were uniquely captured in the benzoyl-chlorine derivatization method. Upset plot of metabolite overlap between five targeted benthic exometabolome studies: Fiore *et al.* 2017 (red), Weber *et al.* 2022 (orange), Garcia *et al.* 2023 - current study (light blue), Weber *et al.* 2022 (blue), and Becker *et al.* 2023 (dark blue). Horizontal bar plots show the set size (sum of metabolites identified for a single study). Vertical bar plots show the intersection size (sum of metabolites unique within or overlapping across studies). Symbol * indicates a study utilizes benzoyl-chloride derivatization method. All other studies utilize a standard PPL-SPE (non-derivatization) extraction approach.

Using the benzoyl-chloride derivatization method, 23 unique metabolites were captured, the highest proportion of unique metabolites of the five investigated studies. Metabolites solely captured using the benzoyl-chloride method consisted of amino acids, amines, pyrimidine nucleosides, and organosulfonic acids; metabolites that have been difficult to capture in the coral exometabolome even with alternative chromatographic and derivatization approaches such as GC-MS.^11^ This approached allowed us to uniquely quantify essential components of central carbon metabolism, carbon fixation in photosynthetic organisms (*e.g.,* aspartate and alanine)^46,47^, and nutrient and substrate transport in prokaryotes (*e.g.,* taurine and putrescine)^48^. The metabolic pathways these compounds comprise are essential for understanding complex organismal processes such as cell viability, virulence, pathogenicity, organism growth and survival. Furthermore, this comparison highlights the large fraction of polar metabolites that have thus far been excluded from DOM chemical investigations, limiting our holistic understanding of chemical exchange and microbial metabolism on coral reefs.

Of the 85 distinct metabolites across the five studies, 40 metabolites were not detected using the derivatization method; 21 of which are intentionally excluded from our targeted list due to (1) successful measurement using PPL-SPE extraction alone thus negating the need for derivatization or (2) lacking the appropriate moieties (amine or alcohol functional groups) for successful benzoyl-chloride derivatization. The 19 metabolites in the BC targeted list that were not detected in this study but were detected in one or more of the SPE studies provide interesting metabolites to query for geographical, ecological, and chemical insights within these reefs. For example, metabolites adenosine, inosine, tryptamine, tyrosine, and xanthosine were detected in all four SPE datasets but fell below the LOD in the benzoyl-chloride derivatized dataset. These metabolites are particularly challenging to detect due to their low standing stock concentrations. All five metabolites were detected in the low fM to pM levels in the SPE-based studies, which utilize much larger extraction volumes (>1L) in comparison to the 25 mL of seawater used for the derivatization protocol. While reduced sample volume allows for increased experimental throughput and replication, which has a multitude of benefits, the limits of detection for these five molecules using the BC method were greater than those determined using PPL-SPE alone. Thus, the set of 19 metabolites undetected using BC derivatization may have been present on our study sites but evaded detection due to their limits of detection. While no single extraction method can capture the DOM pool in its entirety, this comparison demonstrates the complementarity and benefits of BC chemical derivatization relative to more standard approaches.

### Relationship between reef benthos and dissolved metabolites

Ordination-based approaches were used on both habitat and metabolite composition to better understand how biogeographic changes may influence both data types. We used a principal components analysis on benthic composition and found that reefs were significantly structured by both reef and bay (PERMANOVA, Reef: p = 0.001, R^2^ = 0.65, Bay: p = 0.022, R^2^ = 0.14) (Figure S2a). Vectors representing variable contributions of the 11 benthic categories were overlaid on the PCA scores to demonstrate the benthic composition of the reefs. For example, Tektite reef had the highest amount of hard coral and cyanobacterial mats (CYAN), while Ditliff and Cocoloba harbored the highest Substrate (Figure S2a). Joel’s Shoal and Yawzi generally had higher soft coral and turf algae (Figure S2a). Compared to the benthic habitat, the composition of metabolites, as examined with non-metric multidimensional scaling (NMDS) using Bray-Curtis distance, was structured by reef and bay to a lesser extent (PERMANOVA, Reef: p = 0.145, R^2^ = 0.33, Bay: p = 0.082, R^2^ = 0.12) (Figure S2b). Although reef did not significantly structure metabolite composition, it explained 33% of the variance in the metabolite data. Reef habitat components and biogeography are known to influence reef water metabolite composition^22,27,49^, and further sampling across US Virgin Islands reefs in the future is needed to confirm these initial trends.

To understand potential benthic-metabolite relationships, Spearman’s correlations (Figure 5) between metabolite concentrations and benthic proportions were calculated. Correlations illuminated 96 strong (correlation coefficient > 0.7) benthic-metabolite relationships, 48 of which were significant (p < 0.05, SI Excel Table S6). The most prevalent benthic organisms (excluding “other” and substrate) found to have significant correlations include diseased coral, macroalgae, and CCA. This is consistent with previous findings that found benthic primary producers, including CCA, sponges, and algae, variably influenced the DOM pool on coral reefs.^15,28,50,51^ Previous work by Becker *et al.* 2023 further demonstrated the significant impact of disease state on targeted coral exometabolomes.^27^

**FIGURE 5.**
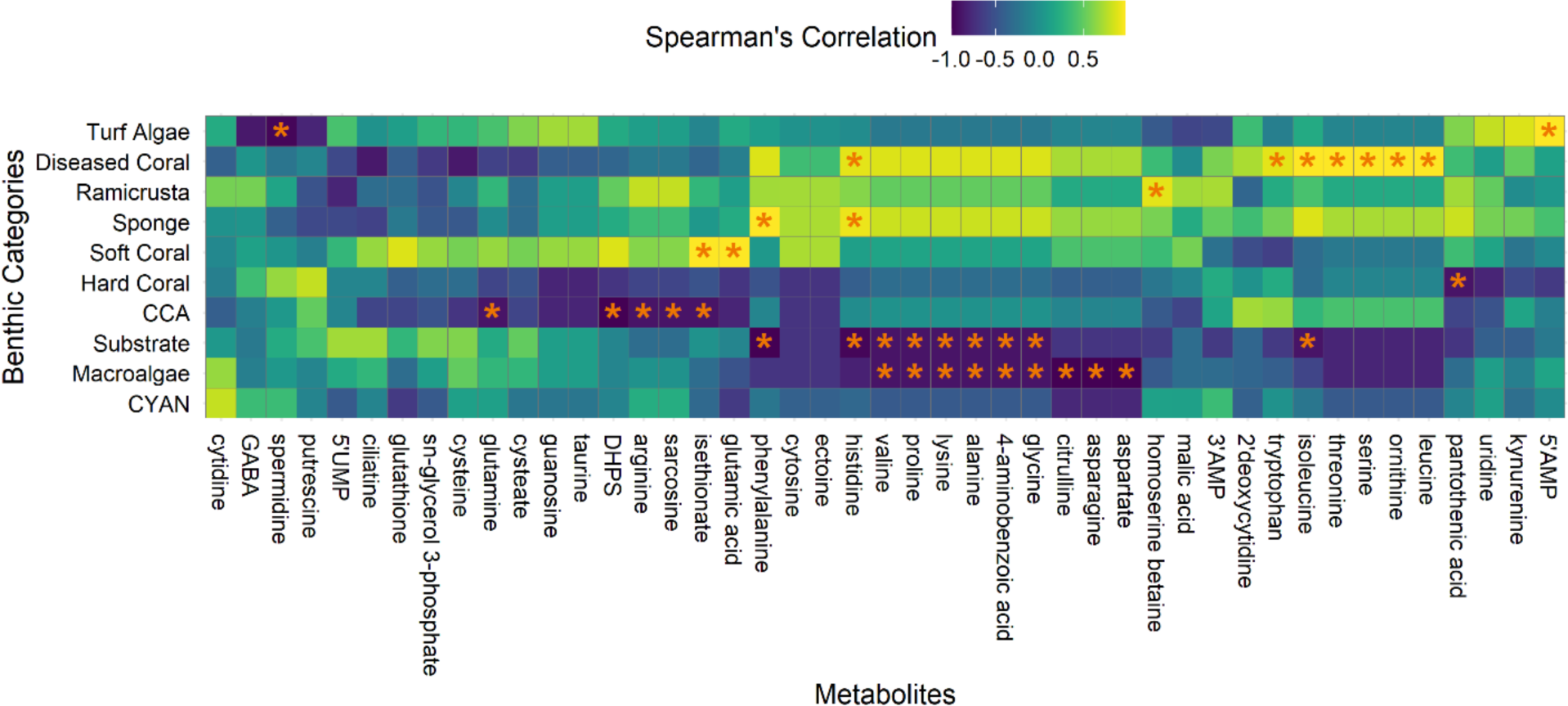
Variations in metabolite concentrations and benthic compositions are strongly related. Spearman correlations between targeted metabolite (x-axis) concentrations and benthic survey data (y-axis). Significant (p < 0.05) correlations are indicated with an orange star. Color corresponds to the strength of the correlation with warmer colors indicating a positive correlation, and cooler colors a negative correlation. CCA = Crustose coralline algae, CYAN = cyanobacterial mats.

Metabolites were tested for significant concentration differences between the five reefs and their respective bays. Due to our small sample sizes when comparing differences between reefs (n = 3-4) a p_adj_ value of 0.1 was selected for our significance threshold. Metabolites homoserine betaine, tryptophan, and γ-aminobutyric acid (GABA) were significantly different based on reef (Figure 6). Tryptophan was found at elevated median concentrations ranging between 49-61 pM at sites JS, YZ, and TK, and fell below the LOD at DL and CO. Interestingly, tryptophan appears to be a core metabolite in the reef exometabolome, as it was also detected in all the prior studies compared earlier. Though the specific role of tryptophan on coral reefs is unknown, bacteria associated with the surface of the coral species *Pocillopora damicornis* and *Acropora aspera* have been shown to exhibit significant levels of chemotaxis towards amino acids, including tryptophan.^52^ There is a need to further investigate the role specific exometabolites play in chemical signaling in coral reef holobiont assemblages.

**FIGURE 6.**
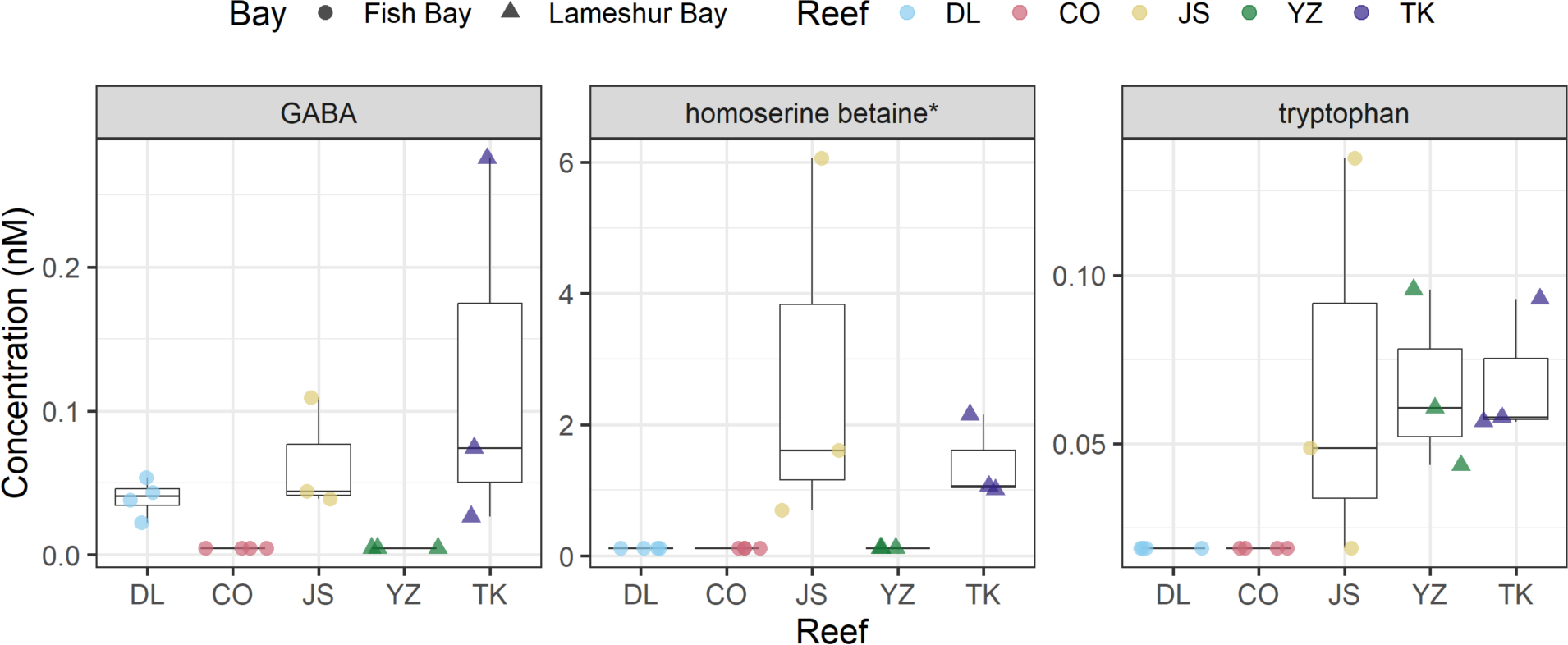
Metabolites GABA, homoserine betaine, and tryptophan show significant differences in concentration across reefs. Boxplots representing the concentrations of significant metabolites by reef (Kruskal-Wallis and Pairwise Wilcoxon Rank Sum Tests, adjusted p-value ≤ 0.1). Boxplots represent the interquartile range (IQR), or the area between the 25 and 75% quantiles, with the median as the line in the center. Lines extend beyond the box to 1.5 × IQR. Points beyond the lines are considered outliers. Color corresponds to each reef, and shape indicates the bay. The x-axis indicates the reef abbreviation and the y-axis displays concentration values (nM). Metabolite names followed by an asterisk (*) indicate metabolites quantified using the BC-derivatization method, but unretained using PPL-SPE alone.

In contrast to tryptophan, metabolites GABA and homoserine betaine have largely gone undetected on coral reefs due to their low (< 1%) PPL extraction efficiencies, with this study being the first report of homoserine betaine in the coral exometabolome. Homoserine betaine is a known compatible solute in the cyanobacterium *Trichodesmium* spp. and has been previously reported in a green alga, *Monostroma nitidium*.^53^ Homoserine betaine was detected at two reefs (JS and TK) and had a significant positive correlation to encrusting red-brown algae *Ramicrusta* (Figure 5), an invasive species on coral reefs shown to overgrow corals and threaten coral recruitment.^54,55^ The presence and relatively high median concentrations of homoserine betaine at JS (1.6 nM) and TK (1.1 nM) is an intriguing finding that requires further investigation due to the limited information available regarding the potential function and presence of homoserine betaine in reef-associated organisms.

GABA is a non-protein amino acid and one of the major inhibitory neurotransmitters in animal central nervous systems. In marine environments, GABA has been shown to play a variety of chemical roles serving as a signaling compound, osmolyte, and carbon substrate.^56–58^ However, the identification and specific role of GABA on coral reefs is not well understood. Weber *et al.* detected GABA in coral reef exudates in the highly protected Jardines de la Reina, Cuba reefs; however, the concentrations fell below their LOQ. We observed elevated median concentrations of GABA ranging from 41-74 pM at sites DL, JS, and TK; in contrast, GABA was absent or fell below our LOD at sites CO and YZ. These findings demonstrate the benefits of chemical derivatization for capturing low abundance metabolites such as GABA that clearly are present within these systems but often evade detection. With the increased replication of our current study, we can confidently quantify GABA on St. John reefs. GABA has been used as an indicator of ecosystem health in snapper,^59^ and has been found as a settlement cue of marine invertebrates.^56^ Coral larvae are also capable of sensing fine-scale settlement cues when they actively explore the benthos in search of highly inducive settlement surface for coral.^60^ GABA did not have any significant correlations to the 11 benthic categories (Figure 5), however, the observed metabolite trends were consistent with elevated total coral coverage (Figure S3). These results suggest GABA could play a chemical cue and thereby increasing hard coral coverage; however additional experiments are needed to specifically address this hypothesis.

While only three metabolites showed significant differences between sites, the metabolites observed reproducibly across sites are also essential to furthering our understanding of marine DOM. For example, DHPS was detected at high concentrations and consistently across reefs. Additionally, DHPS was significantly negatively correlated (Figure 5) with crustose coralline algae (CCA). While this metabolite, and others, were statistically insignificant between sites, their quantification allows for additional hypotheses to be formed and experiments to be conducted investigating the ecological significance of coral exometabolites in relation to varying benthic organisms, and more largely, in the context of marine biogeochemical cycling.

In addition to assessing metabolite variation by reef, we grouped reefs according to location in nearby bay (Fish Bay compared to Lameshur Bay) to evaluate whether coastal hydrodynamics or anthropogenic factors impacted DOM composition. Differences between these two bays that may impact metabolite variation are described in detail in the Supplemental Results. Using a significance threshold of p_adj_ ≤ 0.05, eight metabolites were significantly different between the two bays including cysteate, glutamate, glutamine, glutathione, guanosine, isethionate, sn-glycerol-3-phosphate, and tryptophan (Figure 7). Of these, cysteate, isethionate, and glycerol-3-phosphate have only now been captured in the coral exometabolome using BC derivatization, further highlighting the importance of capturing the polar component of marine DOM. Interestingly, all metabolites, excluding tryptophan, were significantly increased at Fish Bay. These nitrogen-, sulfur-, and phosphorus-rich metabolites are compositionally important in oligotrophic waters where nutrients are limited. However, large influxes or reservoirs of these elements could be a sign of pollution and eutrophication. Specifically, nitrogen enrichment has been shown to have adverse effects on coral growth and calcification,^61^ coral heat resistance,^62^ individual species resilience,^63^ and incidence and severity of coral disease.^64^ TN (inorganic + organic) concentrations were significantly increased (p_adj_ ≤ 0.05) at Fish Bay ([TN]_median_ = 9.15 μM) compared to Lameshur Bay ([TN]_median_ = 6.85 µM), consistent with current hypotheses that increased coastal development may significantly contribute to nitrogen loading (Figure S4). Becker *et al.* 2020 reported inorganic nitrogen and ammonium concentrations from coral reefs (TK, YZ, DL, and CO) located within these two bays, with median concentrations of ∼0.15 μM, suggesting the majority of the TN measurements may be due to organic nitrogen contributions.^65I^ Of the 45 identified metabolites, glutamine and glutamate were found at the highest concentrations across sites, ranging between 744 pM to 50 nM with a median concentration of 2.5 nM (Figure 3). The majority of ammonium in coral reefs is assimilated by the dinoflagellate symbiont via the glutamine synthetase/glutamine:2-oxoglutarate aminotransferase (GS/GOGAT) cycle or by the coral host via GS and/or glutamate dehydrogenase.^66^ Thus, the abundance of glutamine and glutamate could be an indicator of nitrogen assimilation,^66^ with release suggesting symbiont production exceeding host demand, although other organismal uses are also possible. Incorporating exometabolomics approaches into reef monitoring workflows in conjunction with global measurements of TN and TOC enables us to identify what organic compounds are contributing to this bulk pool of essential elements and to what degree in a quantitative manner, furthering our understanding of these complex, rapidly changing marine systems.

**FIGURE 7.**
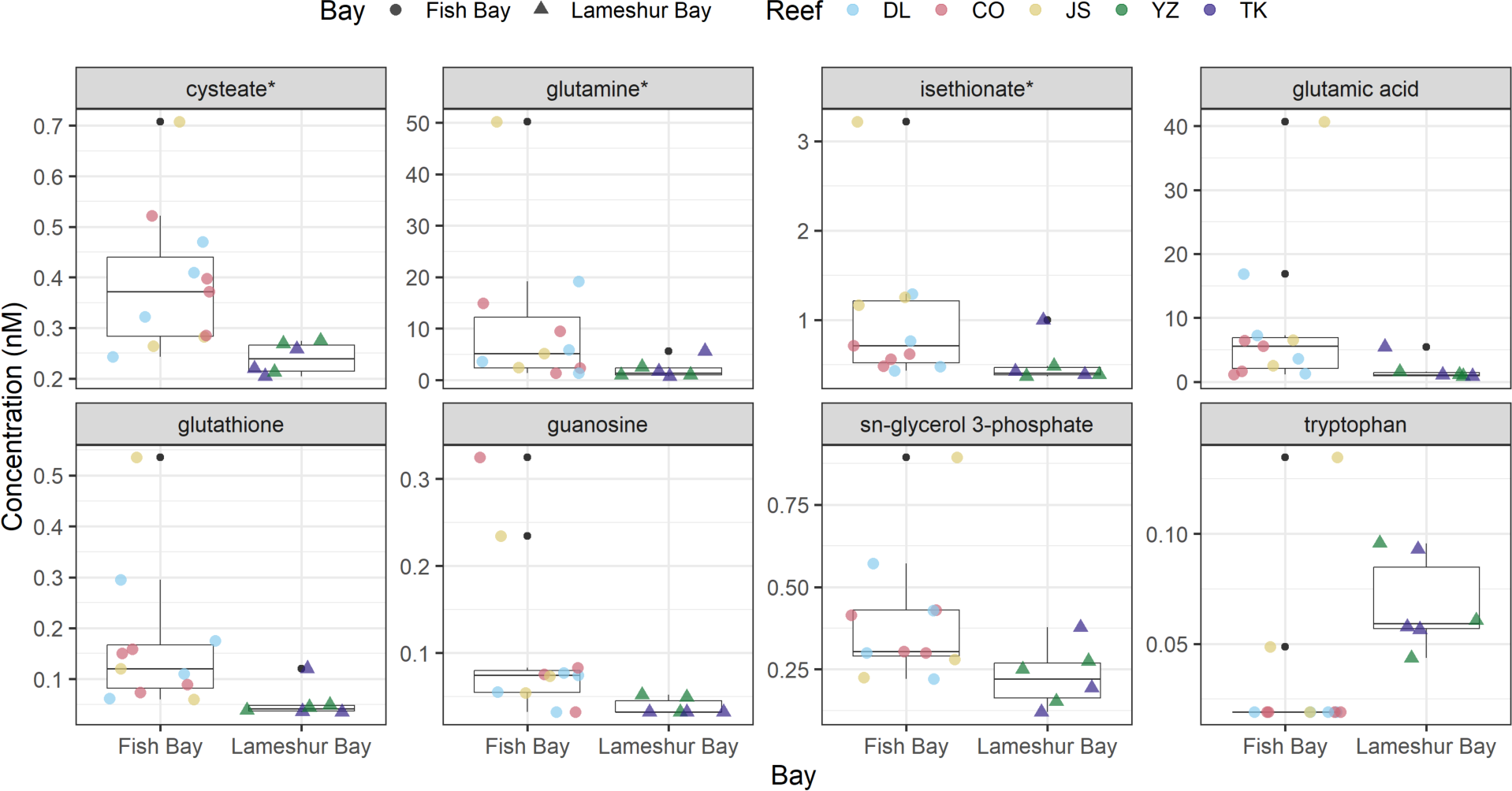
Metabolites significantly elevated at Fish Bay. Boxplots representing the concentrations of significant metabolites on reefs based on bay location (Kruskal-Wallis and Pairwise Wilcoxon Rank Sum Tests, adjusted p-value ≤ 0.05). Boxplots represent the interquartile range (IQR), or the area between the 25 and 75% quantiles, with the median as the line in the center. Lines extend beyond the box to 1.5 × IQR. Points beyond the lines are considered outliers. Color corresponds to each reef, and shapes are indicative of bay. The x-axis labels bay and the y-axis displays concentration values (nM). Metabolite names followed by an asterisk (*) indicate metabolites quantified using the BC-derivatization method, but unretained using PPL-SPE alone.

Here we show that benzoyl chloride derivatization successfully captured 45 polar metabolites, 23 of which were previously undetectable on coral reefs, and that these metabolites were significantly different between individual reefs and across bays. While additional, targeted experiments are needed to relate these observations to their specific source (*i.e.,* runoff, increased biomass, host/symbiont excretion) this data showcases our ability to measure and quantify low abundance, high flux, ecologically relevant metabolites in low volumes of seawater that have previously gone undetected due to methodological limitations. As anthropogenic stressors increase, there is a pressing need to revolutionize our approaches and mindset regarding monitoring strategies. Exometabolomics has the proven potential to elucidate previously overlooked chemical compounds, and novel sample preparation methods, such as chemical derivatization, further increase the suite of molecules we can measure using this approach. As implementation of these methods continues to increase, we will begin to uncover the dynamics of baseline biogeochemical cycles in complex marine environments and harness that information to monitor fragile ecosystems for signs of change related to local and global stressors. Incorporating this method into the standard systems profiling approach (*i.e.,* genomic sequencing, water chemistry, flow cytometry) will allow for additional points of comparison across geographic regions and reef holobiont structure to advance our understanding of the role of labile DOM within complex coral reefs ecosystems, and provide novel targets to investigate their ecological role, sources, sinks, and biomarker potential.

## ASSOCIATED CONTENT

### Supporting Information

The Supporting Information is available free of charge.

### Data Availability

All MS data are under review at MetaboLights^67^ and will be available under accession number MTBLS9008 (https://www.ebi.ac.uk/metabolights/MTBLS9008). A step-by-step benzoyl-chloride derivatization protocol is publicly available on protocols.io (dx.doi.org/10.17504/protocols.io.biukkeuw)^24^. All MATLAB^®^ scripts used for processing the Skyline outputs are available online at https://github.com/KujawinskiLaboratory/SkyMat. R scripts for recreating all figures within the manuscript can be found at https://github.com/bmgarcia/CINAR_Habitat_Metabolomics.

## AUTHOR INFORMATION

### Notes

The authors declare no competing financial interest.

### Author Contributions

- Idea - AA, EBK, LW
- Study design - LW, CCB, AA, EBK, MKS
- Permit - AA
- Sample collection and initial sample processing - AA, CCB, LW
- Sample processing - GS
- Instrument analysis - MKS
- Data processing - BMG, CCB
- Data analysis - BMG, CCB
- Writing - BMG, CCB
- Editing - all authors

## Supporting information

BC_Coral_Exometabolomics_SuppInfo.pdf

BC_Coral_Exometabolomics_SuppInfo.xlsx

## ACKNOWLEDGMENTS

Nadège Aoki and Justin Ossolinski assisted with field sampling and processing, and Krista Longnecker assisted with sample analysis. We thank Erin McParland for helpful discussions regarding experimental design. Samples were collected using National Park Service permit VIIS-2021-SCI-0002. This work was supported by an OCE National Science Foundation award to AA (#1736288) and a NOAA OAR Cooperative Institutes award to AA and EK (#NA19OAR4320074).

## SUPPLEMENTAL INFORMATION

BC_Coral_Exometabolomics_SuppInfo.xlsx

BC_Coral_Exometabolomics_SuppInfo.pdf

