## Supplementary material for "Advancing Reef Monitoring Techniques through Exometabolomics: Quantification of Labile Dissolved Organic Metabolites on Coral Reefs": BC_Coral_Exometabolomics_SuppInfo.pdf

**Table of contents**

1. **Supplemental Methods**
    - Dissolved Organic Carbon
    - Microbial Abundances
    - Benzoyl Chloride Derivatization
    - Targeted Metabolomics UHPLC-MS/MS Gradient and Mass Spectrometer Settings
    - LC-MS Quality Assurance and Control (QA/QC)
    - Metabolite Quantification and Limit of Detection (LOD) Calculations
    - Data Processing QA/QC
2. **Supplemental Results**
    - Differences Between Fish Bay and Lameshur Bay
    - Metabolite Comparison

[1] Department of Soil, Water, and Ecosystem Sciences, University of Florida, Gainesville, FL, United States

- Relationship Between Reef Benthic Composition and Dissolved Metabolites

3. **Supplemental Tables**
   - Table S1. Sampling Site Details and Number Of Metabolites Detected/Quantified
4. **Supplemental Figures**
   - Figure S1. Bubbleplot of Benzoyl-Chloride and SPE Metabolite Comparison
   - Figure S2. NMDS and PCA Ordinations of Benthos (A) and Metabolome (B)
   - Figure S3. Proportion of Total Coral Coverage
   - Figure S4. Flow Cytometry and Inorganic Nutrients
   - Figure S5. Benthic Proportions
   - Figure S6. Ratio Coral to Macroalgae
5. **Supplemental References**

### Supplemental Methods

#### Dissolved Organic Carbon

DOC samples were collected in combusted borosilicate glass vials and acidified with 75 µl concentrated phosphoric acid to pH 2, capped, and stored at room temperature until analysis. NPOC and TN concentrations were analyzed using a Shimadzu TOC-L TOC analyzer[1] with a TNM-L module. Measurements were made using potassium hydrogen phthalate and potassium nitrate as standard solutions.

#### Microbial Abundances

Samples were fixed with 8% paraformaldehyde (1% final concentration), incubated at 4°C in the dark for 20 min, flash frozen via a $LN_2$ dry shipper, and then stored at −80°C prior to analysis. Samples were thawed and stained with Hoechst 34442 (1 µg/mL, final concentration)[2–4] and analyzed using a Beckman-Coulter Altra Flow Cytometer endowed with two argon ion lasers, tuned to UV (200 mW) and 488 nm (1 W) excitation wavelengths. Side and forward scatter as well as fluorescence signals were collected using the appropriate filters designated for Hoechst-bound DNA, phycoerythrin, and chlorophyll. FlowJo software (Tree Star, Inc.) was used to bin populations and estimate the abundances of *Prochlorococcus*, *Synechococcus*, picoeukaryotes, and unpigmented cells.

**Benzoyl Chloride Derivatization**

In batches of 24 samples, 750 µL 8M NaOH was added to each sample and inverted five times to mix. 5 mL of 5% BC in acetone was added to each sample and shaken for five min and then 375 µL of concentrated phosphoric acid ($H_3PO_4$) was added to stop the derivatization reaction. Samples were stored at -20°C until derivatization was complete for all samples. To each sample, 350 µL of $^{13}C$-BC SIL-IS was added, equivalent to a concentration of 5.62 ng of each standard per sample. Samples were dried using a Vacufuge (Eppendorf) until at least 95% of acetone (by weight) was removed from each sample. Samples were then extracted by SPE using 6 mL, 1 g Bond Elut PPL cartridges (Agilent, Santa Clara, CA, USA). PPL cartridges were preconditioned (~6 mL of MeOH followed by ~24 mL of 0.01 M HCl) and samples gravity-loaded. Loaded cartridges were rinsed with four cartridge volumes (~24 mL) of 0.01M HCl, dried for five minutes under vacuum, and eluted with one cartridge volume of MeOH (~6 mL). The eluent was evaporated to near-dryness in a vacufuge, resulting in the formation of a white precipitate. The resulting precipitate was washed with 500 µL 5% MeCN, centrifuged at 1000 *x g* at 22°C for 15 min, transferred to a 2 mL tube, and dried via vacufuge to near-dryness at 30°C. Dried samples were stored at -20°C until LC-MS analysis, at which time each sample was reconstituted in 5% MeCN, transferred to a 2 mL LC vial (with small volume insert), and topped with 5 µL of 100% MeCN. Samples were stored at 4°C until instrumental analysis.

**Targeted Metabolomics UHPLC-MS/MS Gradient and Mass Spectrometer Settings**

The composition of eluent A was MQ water + 0.1% FA, while the composition of eluent B was MeCN + 0.1% FA. The following gradient was used 0-0.5 min 1% B, 0.5-2.0 min 10% B, 2.0-5.0 min 10% B, 5.0-7.0 25% B, 7.0-9.0 25% B, 9.0-12.5 50% B, 12.5-13.0 95% B, 13.0-14.5 95% B, 14.5-14.6 min 1% B, and 14.6-16.0 1% B with a flow rate of 0.5 mL/min and a temperature of 40°C. A diversion valve was used to divert the first 0.8 min of the gradient to waste. MS data were collected in profile mode for both positive and negative ion modes following LC separation. The scan range was 170-1700 m/z. MS1 data was collected at 60,000 resolution (at 200 m/z) with an instrument standard AGC target and a maximum injection time of 50 ms. H-ESI parameters were set as follows: spray voltage +3600V/-2600V, sheath gas - 55 arbitrary units (au), auxiliary gas - 20 au, sweep gas - 1 au, ion transfer tube temperature - 360°C, and vaporizer temperature - 400°C. MS/MS spectra were collected using a targeted list of 79 metabolites at 7,500 resolution (at 200 m/z) with an instrument standard AGC target and a dynamic maximum injection time. The isolation window for MS/MS was set to 1 Da and fragmentation was conducted using higher energy collisional dissociation (HCD) with a normalized collision energy of 35%.

**LC-MS Quality Assurance and Control (QA/QC)**

Four batches of up to 24 samples were needed to derivatize and extract all field samples and generate the (9-point) standard curve. A pooled sample was made by combining 5 µL aliquots from all derivatized field samples across all four batches. The pooled sample was used to condition the UHPLC column before executing the sample queue and was repeatedly injected every nine samples to assess instrument drift and for downstream data processing. MQ water blanks were included at the beginning and end of each queue to assess sample carry-over and LC-MS system contamination. A derivatized matrix blank was generated using seawater collected ~1 mile offshore (18 17.127°N, 064 44.312°W, 31.6 m sampling depth) to assess (1) the baseline concentration of non-reef dissolved-phase metabolites present in seawater after derivatization, and (2) contaminants from sample handling, derivatization, and the extraction process. Data for the standard curve was collected at the beginning of the queue in order of ascending concentration, followed by field samples in a randomized order. Samples were reinjected for positive and negative ionization modes, and run order was kept consistent across modes.

**Metabolite Quantification and Limit of Detection (LOD) Calculations**

Metabolites were quantified by calculating the light-to-heavy peak area ratio of each precursor ion and then converted to ng added concentrations based on the linear regression. Concentrations in ng added were then converted to nM values using a MATLAB® script (*convertMoles.m*). The concentration of metabolites in each sample was calculated by dividing the ng added concentration by the volume of filtrate (25 mL) that was derivatized and subsequently passed through each PPL column and multiplying by the molecular weight of each respective metabolite standard. Limit of detection (LOD) and quantification (LOQ) were determined for each metabolite as shown in Equations 1 and 2; where $s_b$ is the standard deviation of the y-intercept and $m$ is the slope of the calibration curve.[5] Samples with a metabolite concentration less than the corresponding LOD were replaced with 1/10th the calculated LOD value for downstream statistics.

$$LOD = 3.3 * \frac{s_b}{m} \qquad \text{Equation 1}$$

$$LOQ = 10 * \frac{s_b}{m} \qquad \text{Equation 2}$$

**Data Processing QA/QC:**

Metabolites were initially filtered out by manual inspection of integrated peaks in Skyline. Samples were collected in triplicate, and metabolites present in less than 2/3 of the replicates were replaced with 1/10th the LOD value of the corresponding metabolite. Metabolites with a coefficient of variation (CV) < 10% across all samples were removed. Metabolites present in (1) both polarity modes, (2) multiple derivatization states (1-3 benzoyl ring additions), or (3) multiple adduct ionization states were assessed for intensity, linearity, and presence/absence retaining only the highest performing metabolite for each set of duplicates. Post QA/QC, a total of 51 unique metabolites (24 quantified in positive mode and 27 quantified in negative mode) were retained. Metabolites with a standard deviation of zero were removed resulting in a total of 45 detected metabolites. Sample outliers were determined on a 'by reef' basis by calculating the pairwise Euclidean distance between each sample within a reef. A sample was considered an outlier if the average Euclidean distance of that sample did not fall within 1.5 times the interquartile range +/- the upper (75th) and lower (25th) quartile - the equivalent of +/- 2.698σ. No sample outliers were found. For metabolite concentrations that exceeded the 9-point standard curve, the linearity of the regression up to the detected concentration was confirmed by generating an extended standard curve encompassing the detected concentration ranges. All detected metabolites were linear beyond 100 ng added thus allowing quantification of all metabolites. The initial standard curve was used for quantification in lieu of the extended curve to eliminate any batch effects between the study samples and the externally generated extended curve generated at a later date.

### Supplemental Results:

#### Differences Between Fish Bay and Lameshur Bay

Reefs DL, CO and JS in Fish Bay, and TK and YZ are in Lameshur Bay. Both bays have inputs from mangrove forests. Fish Bay's coastline is highly developed, harbors one of the largest watersheds on St. John, and is considered impacted by human development due to erosion and sedimentation from development and bay contamination from septic systems, household pollutants, and pesticides.[6,7] While Lameshur Bay has a watershed area of similar size to that in Fish Bay, the number of developments and roads at Lameshur Bay are substantially lower.[6] Additionally, JS which falls on the eastern edge of Fish Bay may be exposed to increased water cycling and less particle retention. CCA, diseased coral, and soft coral were significantly different

($p_{adj} \leq 0.05$) between the two bays, with higher proportions of CCA and diseased coral in Lameshur Bay and higher proportions of soft coral at Fish Bay. Land-based sources of pollution (*e.g.,* coastal development and agricultural runoff) are a major factor in coral reef degradation. However, the amounts and types of pollutants present on coral reefs are not well characterized, and less is known about the effects of these contaminants on coral health. Contextualizing differences between sampling locations and potential natural and anthropogenic inputs is essential to furthering our understanding of these relationships.

**Metabolite Comparison**

Six metabolites were found in all sample sets regardless of geography, sample preparation method, or experimental approach (Figure S1). These metabolites included amino acids (kynurenine, tryptophan, phenylalanine), B-vitamin pantothenic acid, purine nucleoside guanosine, and benzoic acid 4-aminobenzoic acid. These metabolites were present in protected regions such as Jardines de la Reina, and throughout the Caribbean and Florida coast, areas overcome with incidents of coral disease and bleaching. Metabolites kynurenine, pantothenic acid, guanosine, and 4-aminobenzoic acid have been reported in aged oligotrophic seawater from the Bermuda Atlantic Time-series Study (BATS) at 1m (shallower, but comparable to the reef depths investigated within these studies).[8] Metabolite concentrations for these four compounds were similar between the current study and those reported by Widner *et al.*[8], indicating these metabolites could be less associated with the reef benthos and more likely recurring members of the overall seawater metabolome. In contrast, tryptophan and phenylalanine were not detected in the open ocean samples at 1m reported by Widner *et al.*[8], but have been reported in off-reef surface water samples by Weber *et al.* 2020.[9] Corals can synthesize amino acids *de novo*,[10] including eight essential amino acids: valine, isoleucine, leucine, tyrosine, phenylalanine, histidine, methionine, and lysine, and have previously been shown to release amino acids,[9,11–14] vitamins,[9] and nucleosides[9,14,15] into the environment. Tryptophan and phenylalanine are synthesized from intermediates in glycolysis and the pentose phosphate shunt;[10] genes involved in the pentose-phosphate pathway have been positively correlated with the algal cover on microbialized reefs.[16] Thus, the recurring presence of amino acids phenylalanine and tryptophan on coral reefs may be related to microbialization, however, further monitoring and investigations into the sources of these metabolites is necessary.

Metabolites detected in multiple studies but not uniformly across all five present the interesting question of metabolite role, geographic and ecological influences, and differences in experimental design. The largest set of overlapping metabolites was found between Weber *et al.* 2020 and the current study, both field benthic seawater collections from Cuba and the USVI, respectively, sharing nine common metabolites. Additionally, Weber *et al.* 2020, Weber *et al.* 2022, and Becker *et al.* 2023 also shared nine overlapping metabolite identifications. Determining definitive correlations between these variable metabolite observations is challenging given the low number of targeted exometabolomic studies available for comparison. These findings further highlight the importance of elucidating the composition of marine DOM and incorporating exometabolomics into future coral reef monitoring experimental design strategies, as highlighted in a recent perspective by Apprill *et al.*[17]

**Relationship Between Reef Benthic Composition and Dissolved Metabolites**

To discern the potential sources and sinks of dissolved phase metabolites, the benthic composition of each reef was determined. Hard coral, macroalgae, and turf algae were the most abundant categories across all reefs, whereas crustose coralline algae (CCA) and diseased coral had the smallest overall proportions (Figure S5). Benthic composition varied significantly across reefs (SI Excel Sheet S3). Non-parametric Kruskal-Wallis tests revealed significant differences ($p < 0.05$) between reefs for cyanobacterial mats, diseased coral, hard coral, macroalgae, *Ramicrusta*, soft coral, sponge, and turf algae. Pairwise comparisons using the Wilcoxon rank sum test with Benjamini & Hochberg correction showed significant ($p_{adj} \leq 0.1$) pairwise differences for cyanobacterial mats, diseased coral, macroalgae, soft coral, and turf algae.

Additionally, we compared the total coral coverage (summation of hard and soft corals) across sites (Figure S3). CO and YZ had low coral coverage (median proportion < 30%), whereas DL, JS, and TK had higher coral coverage (median proportion values > 30%). The hard coral to macroalgae ratio (Figure S6) was also investigated given the large contribution of macroalgae to the PC1 and the known impact of macroalgae on coral reef DOM (Figure S2).[18] Increased algal biomass produces excess photosynthate that is released into the water in the form of DOC.[16] The addition of large influxes of DOC into typically oligotrophic reefs supports microbialization which can lead to pathogenesis or anoxic conditions for the coral host and their symbionts.[16] DL and TK had roughly equal proportions of hard coral and macroalgae, whereas YZ and JS had twice as much hard coral compared to macroalgae. CO was the only site to have more macroalgae than hard coral. These trends were similar to those observed when visualizing the reefs using PCA (Figure S2) suggesting these high contributing variables are representative of the differences we observe between reefs.

### Supplemental Tables

TABLE S1. **Sampling site details and number of metabolites detected/quantified.** Quantified metabolite numbers represent the number of detected metabolites at each reef that had at least one measured value greater than the calculated LOD.

**Total # detected metabolites = 45**

| **Reef (Abbreviation)** | Bay | Latitude | Longitude | Reef Depth (m) | Sample Size (n) | Detected metabolites | Quantified metabolites |
|---|---|---|---|---|---|---|---|
| Ditliff (DL) | Fish | 18.313 | -64.764 | 6.10 | 4 | 35 | 33 |
| Cocoloba (CO) | Fish | 18.315 | -64.761 | 7.01 | 4 | 35 | 29 |
| Joel’s Shoal (JS) | Fish | 18.313 | -64.757 | 8.8 | 3 | 44 | 42 |
| Tektite (TK) | Lameshur | 18.309 | -64.723 | 8.3 | 3 | 39 | 36 |
| Yawzi (YZ) | Lameshur | 18.314 | -64.726 | 8.4 | 3 | 39 | 28 |

### Supplemental Figures

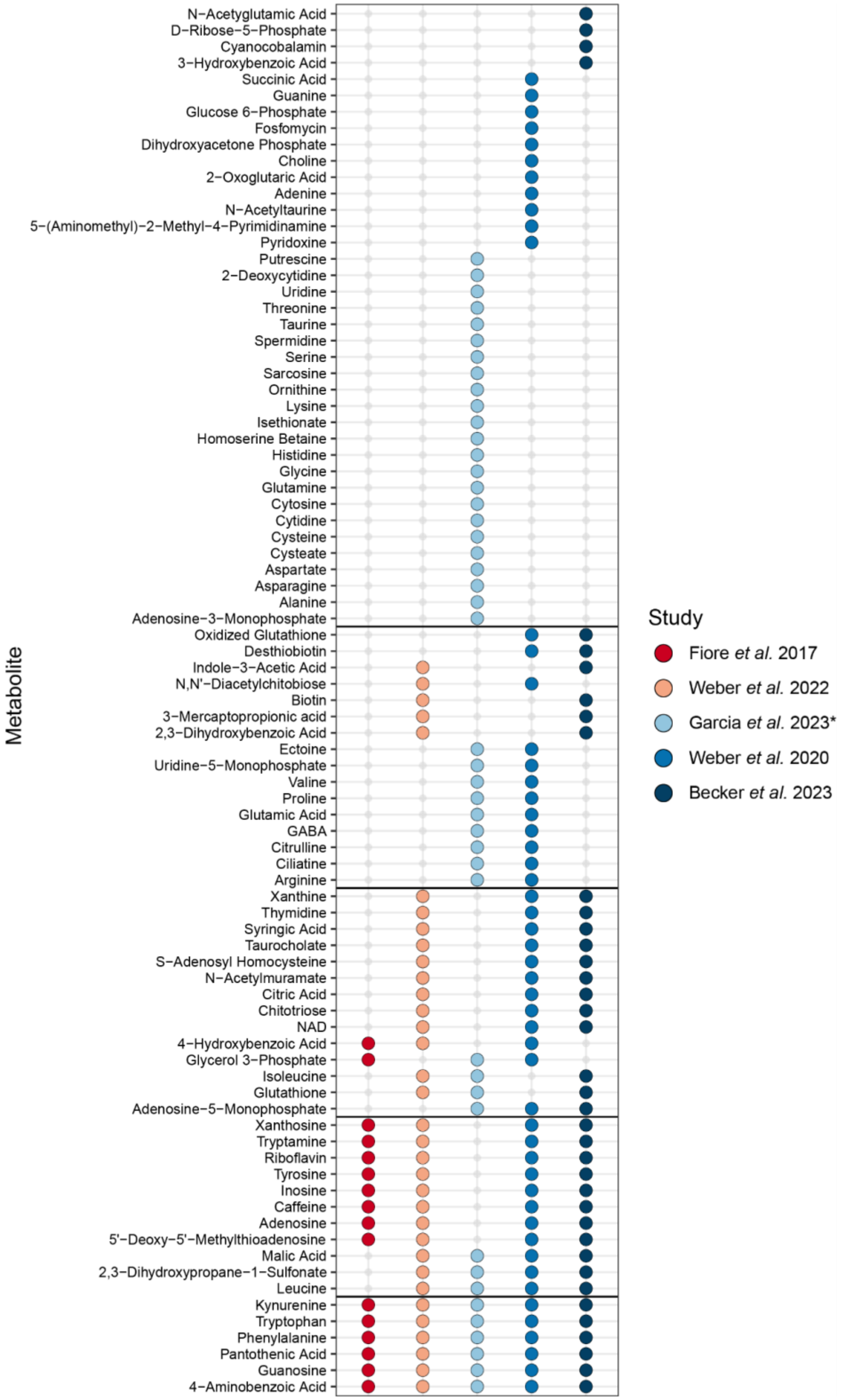

FIGURE S1. **Bubbleplot showing the identified metabolite comparison across targeted coral exometabolomics datasets.** Metabolites are depicted on the y-axis and grouped based on the number of studies (1-5) that metabolite was identified in, with the most unique metabolites at the top and the metabolites with the most overlap at the bottom. Colors correspond to the study the metabolite was found in, consistent with those used in Figure 4. The absence of a bubble indicates the metabolite was not identified in that corresponding dataset.

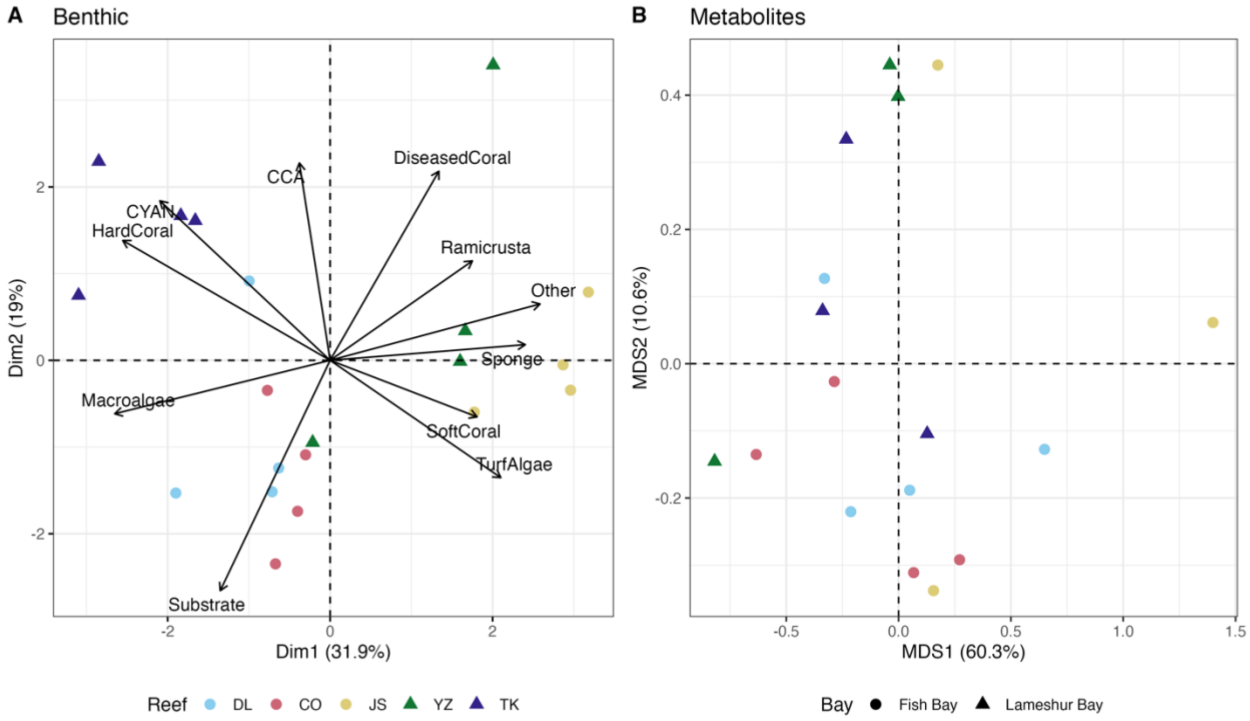


FIGURE S2. **NMDS and PCA Ordinations of Benthos and Metabolome.** (A) Bray-Curtis dissimilarity between targeted metabolomics samples represented by NMDS. (B) Principal Components Analysis (PCA) of benthic survey data. Color is indicative of the reef.

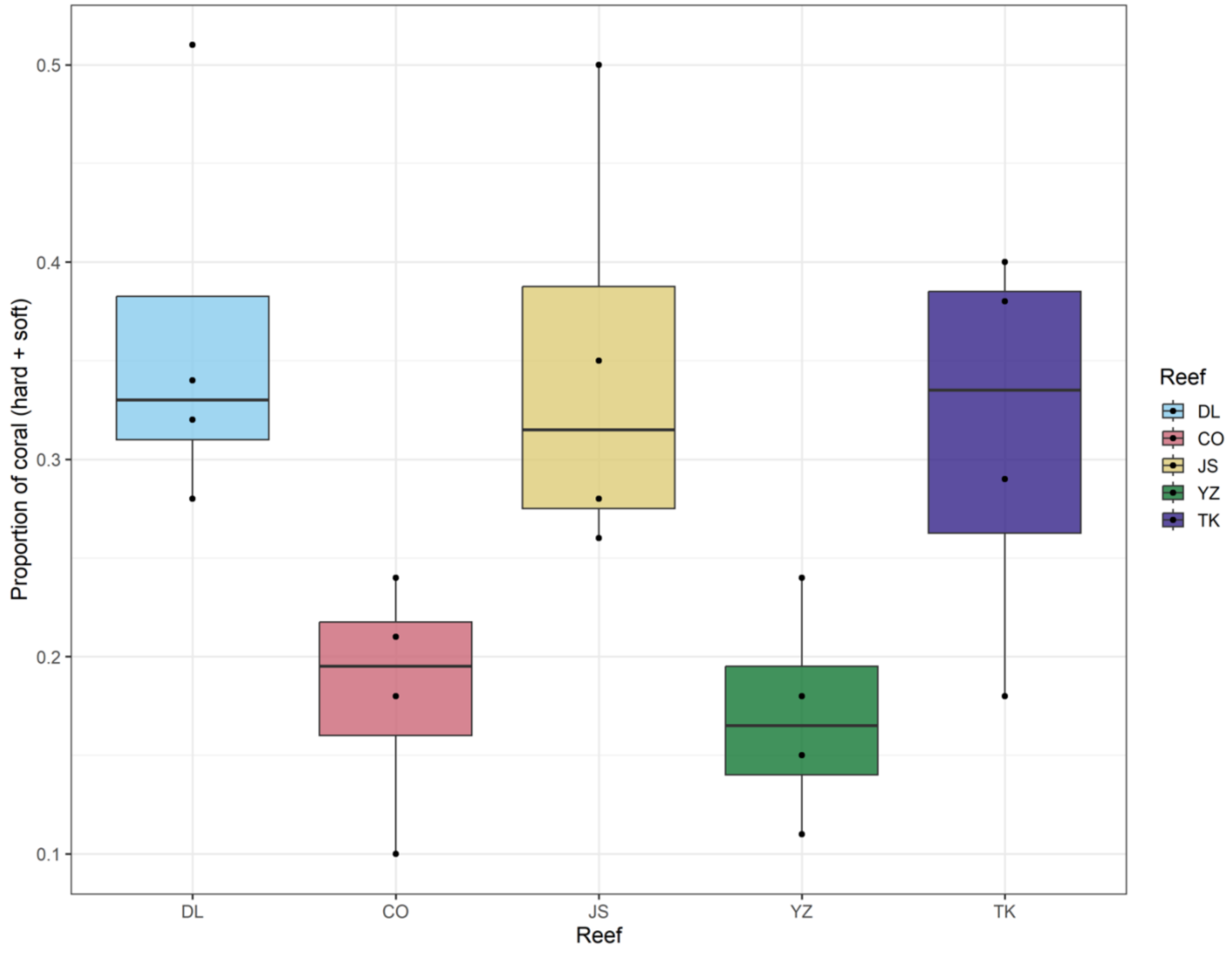


FIGURE S3. **Proportion of coral (hard + soft) calculated for the benthic surveys of each sampling site.** Each datapoint represents a survey transect and each color represents a unique sampling site. Boxplots represent the interquartile range (IQR), or the area between the 25 and 75% quantiles with the median as the line in the center. Lines extend beyond the box to 1.5 × IQR. Points beyond the lines are outliers.

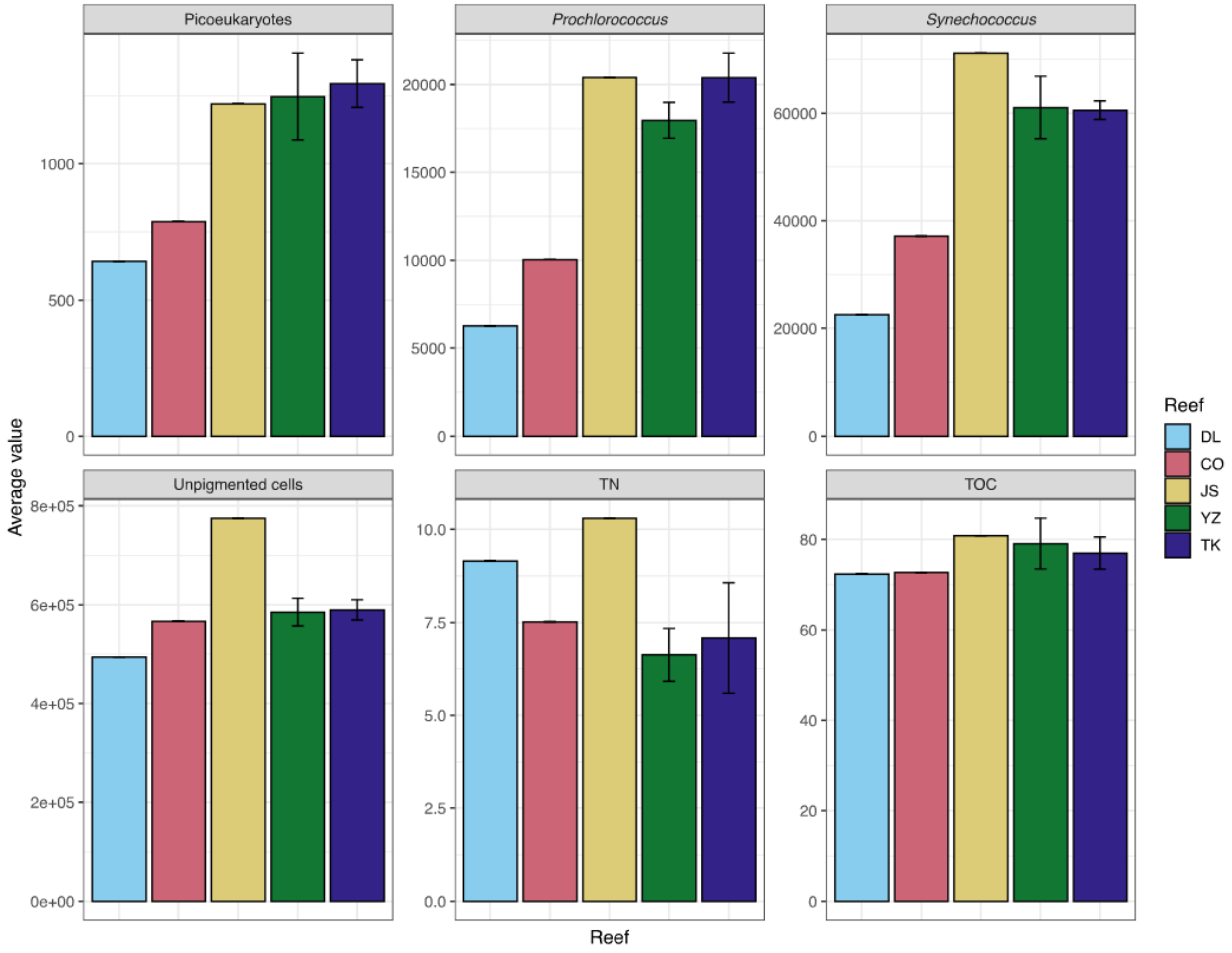


FIGURE S4. **Cell abundances from flow cytometry and organic and inorganic nutrient concentrations at each site.** Bar plot of (a) Picoeukaryotes, (b) *Prochlorococcus*, (c) *Synechococcus*, (d) heterotrophic (unpigmented) bacteria and archaea, (e) total nitrogen (TN - inorganic + organic), and (f). Bar heights represent the average value. Sites with error bars indicate one standard deviation from the mean. Reefs without error bars had only a single measurement to report.

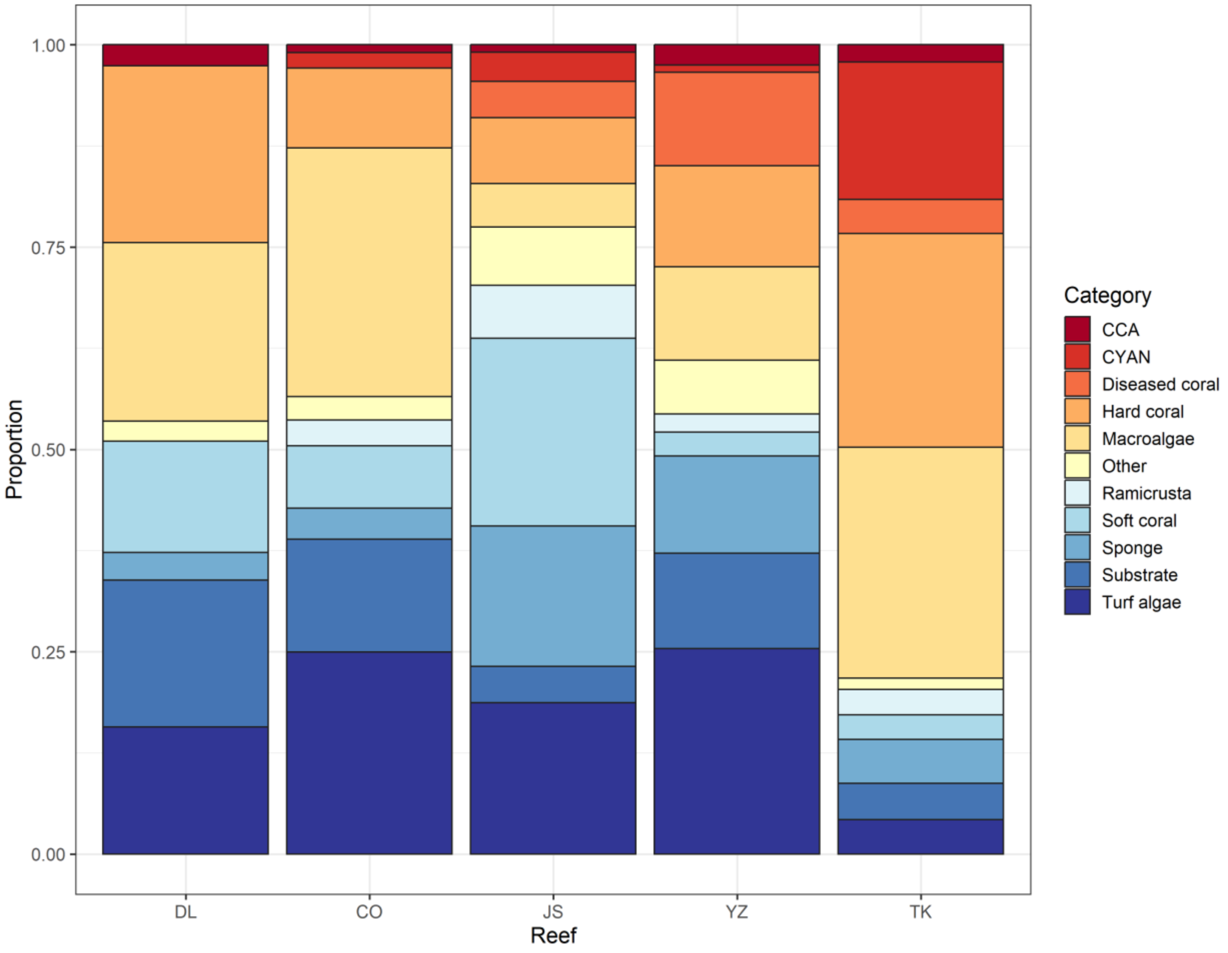


FIGURE S5. **Benthic composition.** Stacked barplot representing the benthic composition at each site calculated using the point intercept benthic surveys. Larger areas indicate a higher proportion of that specific benthic category at the corresponding site. Total proportions add up to one or 100%. Color is representative of the eleven benthic categories used in the benthic survey.

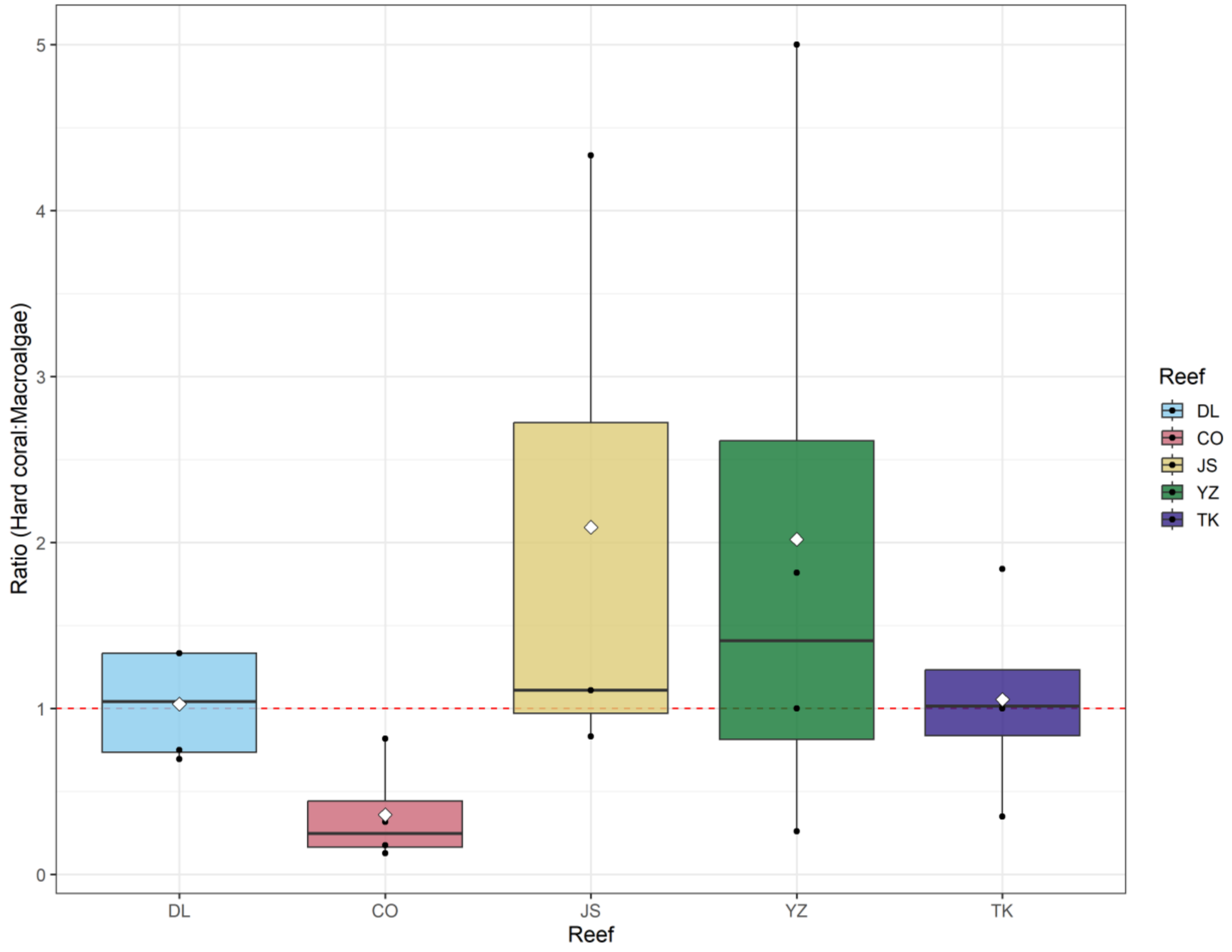


FIGURE S6. **Ratio of hard coral to macroalgae calculated for the benthic surveys of each sampling site.** The red dashed line represents a 1:1 ratio or equal proportions of hard coral and macroalgae. Each datapoint represents a survey transect and each color represents a unique sampling site. Boxplots represent the interquartile range (IQR), or the area between the 25 and 75% quantiles with the median as the line in the center. Lines extend beyond the box to 1.5 × IQR. Points beyond the lines are outliers. A white diamond was used to represent the average value for each site.

(18) Nelson, C. E.; Goldberg, S. J.; Wegley Kelly, L.; Haas, A. F.; Smith, J. E.; Rohwer, F.; Carlson, C. A. Coral and Macroalgal Exudates Vary in Neutral Sugar Composition and

Differentially Enrich Reef Bacterioplankton Lineages. *ISME J.* **2013**, *7* (5), 962–979. https://doi.org/10.1038/ismej.2012.161.
